# Near-Atomic Resolution Structure Determination in Over-Focus with Volta Phase Plate by Cs-corrected Cryo-EM

**DOI:** 10.1101/148205

**Authors:** Xiao Fan, Lingyun Zhao, Chuan Liu, Jin-Can Zhang, Kelong Fan, Xiyun Yan, Hai-Lin Peng, Jianlin Lei, Hong-Wei Wang

## Abstract

Volta phase plate (VPP) is a recently developed transmission electron microscope (TEM) apparatus that can significantly enhance the image contrast of biological samples in cryo-electron microscopy (cryo-EM) therefore impose the possibility to solve structures of relatively small macromolecules at high resolution. In this work, we performed theoretical analysis and found that using phase plate on objective lens spherical aberration (Cs)-corrected TEM may gain some interesting optical properties, including the over-focus imaging of macromolecules. We subsequently evaluated the imaging strategy of frozen-hydrated apo-ferritin with VPP on a Cs-corrected TEM and obtained the structure of apo-ferritin at near atomic resolution from both under- and over-focused dataset, illustrating the feasibility and new potential of combining VPP with Cs-corrected TEM for high resolution cryo-EM.

**Highlights:** The successful combination of volta phase plate and Cs-corrector in single particle cryo-EM.

Near-atomic structure determined from over-focused images by cryo-EM. VPP-Cs-corrector coupled EM provides interesting optical properties.

**In Brief:** We took the unique advantage of the optical system by combining the volta phase plate and Cs-corrector in a modern TEM to collect high resolution micrographs of frozen-hydrated apo-ferritin in over-focus imaging conditions and determined the structure of apo-ferritin at 3.0 Angstrom resolution.

## Introduction

The recent technical breakthroughs, such as the invention of direct electron detector (Faruqi and Henderson, 2007, McMullan et al., 2014) and the implementation of new image processing algorithms (Brilot et al., 2012, Li et al., 2013, Scheres, 2012), have brought single particle cryo-electron microscopy (cryo-EM) a stunning capability to determine the structure of macromolecules at atomic or near-atomic resolution (Bai et al., 2015, Cheng et al., Nogales and Scheres, 2015). As the central instrument of the technology, a modern transmission electron microscope (TEM) has a rather complex electron optics that still has a lot of space to be further improved. The community of material science has witnessed the application of a few new electron optical apparatuses to reveal structures at sub-atomic resolution. It is tempting to explore if the cryo-EM technology can benefit from new optical-tuning equipment, including the phase plate and Cs-corrector.

An under-focus data collection strategy on conventional TEM (CTEM) has been exploited for decades according to the contrast transfer theory that predicts a sine contrast transfer function (CTF) with a small percentage of amplitude contrast for weak-phase biological specimens (Frank, 2006). Adding a phase plate at the back-focal plane of the objective lens could modulate the CTF as a cosine function (Zernike, 1942, Danev and Nagayama, 2001). A major advantage of using the phase plate is the strong boost of image contrast because more low frequency information of the structure is maintained in the image, which is of particular usage to biological specimens (Glaeser, 2013). Unwin used spider thread to experimentally demonstrate its potential usage in 1970s (Unwin, 1973). A practically usable carbon-film-based Zernike phase plate was implemented by Nagayama and Danev (Danev and Nagayama, 2001). Around the same time, various designs of phase plate were also proposed and tested (Danev et al., 2009, Lentzen, 2004, Hosokawa et al., 2005). Using the carbon-film-based Zernike phase plate, a few single particle structures have been obtained using cryo-EM (Dai et al., 2014, Taylor et al., 2013, Danev and Nagayama, 2001, Danev and Nagayama, 2008, Danev et al., 2009, Nagayama, 2008). It has also been used in cryo-electron tomography to study cellular structures (Danev et al., 2010, Murata et al., 2010, Hosogi et al., 2011, Fukuda and Nagayama, 2012, Guerrero-Ferreira and Wright, 2014). In 2015, a new type of phase plate using a voltage-charged film, so called volta phase plate (VPP), was shown to be able to give high contrasted images of biological molecules with much less fringes than the Zernike phase plate (Danev et al., 2014). In combination with the direct electron detector, VPP allowed the reconstruction of the 20S proteasome complex at 3.2 Angstrom resolution from in-focus images (Danev and Baumeister, 2016).

Ideally, a TEM equipped with a phase plate is able to faithfully record the structural information of an object at all frequencies when the specimen is in-focus of the objective lens. The presence of spherical aberration in the objective lens, however, causes the information distortion in the images especially at high frequency. Such a problem may be overcome by the usage of a Cs-corrector of the objective lens. Cscorrector-equipped microscope has been shown to be able to solve the structure of ribosomes at near atomic resolution (Fischer et al., 2015, Fischer et al., 2016). No attempt has yet been reported to use the combination of phase plate and Cs-corrector for high resolution cryo-EM, although theoretical analysis and simulation suggests that a combination would maintain more structural information at high spatial frequencies (Evans et al., 2008, Gamm et al., 2008).

In this work, we performed theoretical analysis and noted that the combination of a phase plate and Cs-corrector can generate images suitable for high resolution structural determination in both the under- and over-focus imaging conditions. Using a VPP on a Cs-corrector-equipped electron microscope, we imaged ice-embedded apoferritin molecules under either under- or over-focus imaging conditions. Single particle analysis of both the under- and over-focused datasets led to reconstructions at ~3.0 Angstrom resolution. The two datasets can also be combined and treated as a single dataset reconstructed at 3.0 Angstrom resolution, illustrating a novel imaging capability for high resolution cryo-EM by the combination of VPP and Cs-corrector.

## Results

### CTF of under- and over-focus cryo-EM with phase plate and Cs-corrector

For thin biological specimens in TEM, the weak-phase approximation predicts that the Fourier transform of a TEM image as a multiplication of the Fourier transform of the object’s projection and the CTF of the TEM’s electron-optic system (Frank, 2006). A simplified form of the CTF can be written as:
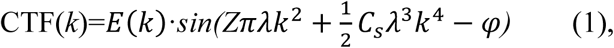

where *k* is the spatial frequency in Fourier space; *λ* is the electron wave length with given acceleration voltage; *Z* is the defocus value with positive sign for over-focus and negative sign for under-focus imaging condition; *C_s_* is the spherical aberration of the TEM objective lens; *φ* is an integrated phase shift combining the amplitude contrast and phase plate effects; and *E(k)* is the envelope function that modulates the amplitude of CTF. We can calculate the CTFs in respective of the spatial frequency under different imaging scenarios using Equation 1.

For CTEM without phase plate, the *φ* is a rather small positive value contributed only by the amplitude contrast. When the Z is positive (over-focus imaging condition), the CTF starts with a small negative value at zero frequency and crosses the frequency axis rapidly at low frequency (Figure 1A, CTEM blue curve). This generates a complicated contrast deviation for low resolution information and makes it rather difficult to determine and correct CTF accurately in real images. In contrast, the under-focus imaging condition (negative Z value) makes the CTF with a broader bandwidth (from the origin to the first zero) in low frequency area (Figure 1A, CTEM red curve) and allows convenient CTF determination and correction so has been the norm for cryoEM imaging for CTEM. The same phenomenon holds for cryo-EM on a Cs-corrected EM (Figure 1A, Cs-corrector only). As predicted by the CTFs, under-focused and over-focused images show opposite specimen contrasts (Figure 1B).

**Figure 1.**
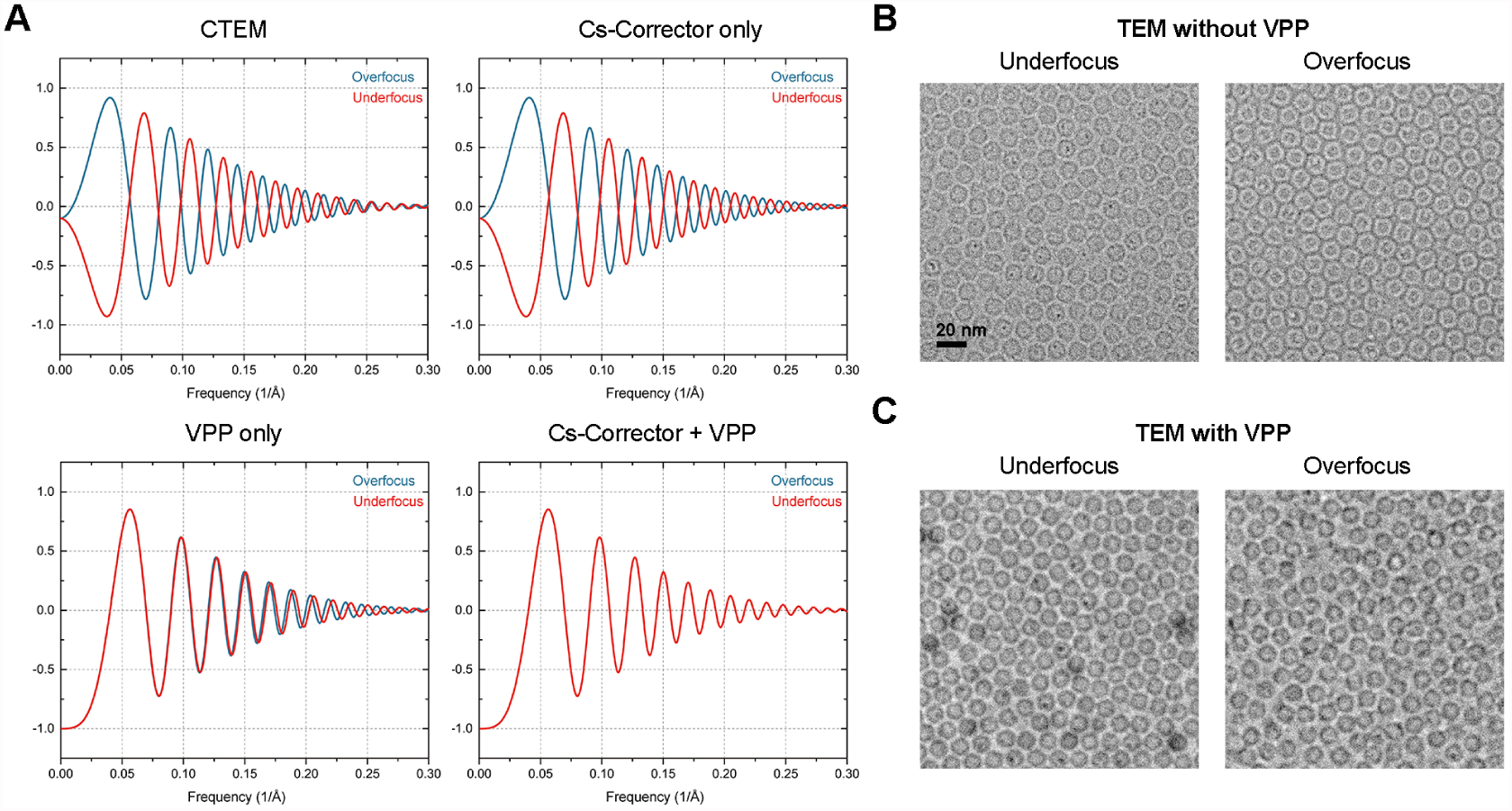
CTF and cryo-EM images by VPP-Cs-corrector coupled EM. (A) Simulated contrast transfer functions (CTF) in different scenarios. For conventional TEM (CTEM), the spherical aberration (Cs) is 2.7 mm with ~10% amplitude contrast. For Cs-corrected TEM, Cs is set to 0.001 mm (1μm). An ideal VPP which introduces a 7c/2 phase shift is applied in this simulation. −0.3 and +0.3 μm defocus values are used for under- or over-focus imaging conditions in all simulations, respectively. (B) Micrographs of apo-ferritin particles collected with −2.0 μm under-focus (left) and +2.0 μm over-focus (right) imaging conditions using Cs-corrected TEM. (C) Micrographs of apo-ferritin particles collected with −0.5 μm under-focus (left) and +0.5 μm over-focus (right) imaging conditions using Cs-corrected TEM with VPP.

When applying a phase plate that can generate a phase shift of ~90 degrees, the *φ* value is close to *π*/2 and the CTF becomes a cosine-like function (Figure 1A). As a result, there is no flip of sign of CTF at the very low frequency irrespective of the Z value’s sign, therefore both under- and over-focused imaging conditions lead to the same image contrast (Figure 1C). In principle, for both imaging conditions, the CTF parameters, including the Z and *φ*, can be accurately determined by fitting Equation 1 in the power spectrum of a raw image with the known Cs value. The CTF can subsequently be corrected for image processing with the estimated parameters. We noted that the relatively large Cs value on a normal TEM, i.e. 2.7 mm of Cs on a Titan Krios, causes significant difference of CTF curves for under- and over-focused images especially at the high frequency range (Figure 1A, VPP only). This means that for phase-plate images, a CTF fitting program must take care of the under- or over-focus imaging conditions separately whereas the sign of Z value cannot be directly obtained from either the raw image or the power spectrum. As far as we know, the currently available CTF fitting programs do not have the capability to meet such a need. This problem, however, can be eliminated by drastically reducing the Cs value in a TEM equipped with a Cs-corrector. With a very small Cs value (~0.001 mm), the second term for the sine function in Equation 1 can be neglected. The equation thus becomes:
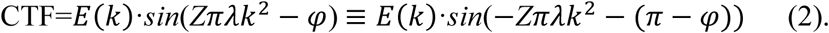

If *φ* is modulated to *π*/2, the above equation predicts that a same CTF curve would be generated by under- and over-focus imaging conditions with the same absolute defocus value (Figure 1A, Cs-corrector plus VPP). As a matter of fact, as long as the *φ* is around *π*/2 (not necessarily to be exact), we could treat the over-focused images (positive Z value) in the same way as the under-focused images (negative Z value) except that we need to use a *φ*′ = (*π* — *φ*) to perform the CTF fitting. The most current version of CTFFIND 4.15 (Rohou and Grigorieff, 2015) and Gctf-1.06 (Zhang, 2016) programs can be used to search for both Z and *φ* values for an optimal fitting. The estimated parameters can be directly used for CTF correction in later image processing steps. In conclusion, the combination of a Cs-corrector and a phase plate allows to image the biological cryo-EM specimens using under- or over-focus imaging conditions and treat the images in exactly the same way.

### Single particle reconstruction of apo-ferritin by VPPCs-corrector coupled cryo-EM

To test the above theory, we established a procedure (Figure S1 and Method) to collect cryo-EM images of apo-ferritin embedded in vitreous ice using the VPP-Cs-corrector coupled cryo-EM. We intentionally collected under- and over-focused datasets of the specimen with the phase shift ranging from 0.2-0.8π. Three datasets were thus generated: 1) −0.4 to −1.0 μm under-focused dataset, 2) 0.4 to 1.0 μm over-focused dataset, and 3) the mixture of under-focused (dataset 1) and over-focused (dataset 2) images. We treated all the 3 datasets as under-focused images for CTF parameter determination and single particle analysis following the standard Relion image processing strategy (Kimanius et al., 2016). We were able to calculate the 2D class averages that showed fine details of the apo-ferritin in different views indistinguishable among the three datasets (Figure 2A). For all three datasets, we successfully performed 3D classification and refinement and obtained 3D reconstructions of apo-ferritin at 2.9 Å (under-focused dataset, Figure S2A), 3.2 Å (over-focused dataset, Figure S2B) and 3.0 Å (mixed under-/over-focused dataset, Figure 2B and S2C), respectively. The three reconstructions appeared almost exactly the same. The subtle resolution difference seems related to the dataset quality.

**Figure 2.**
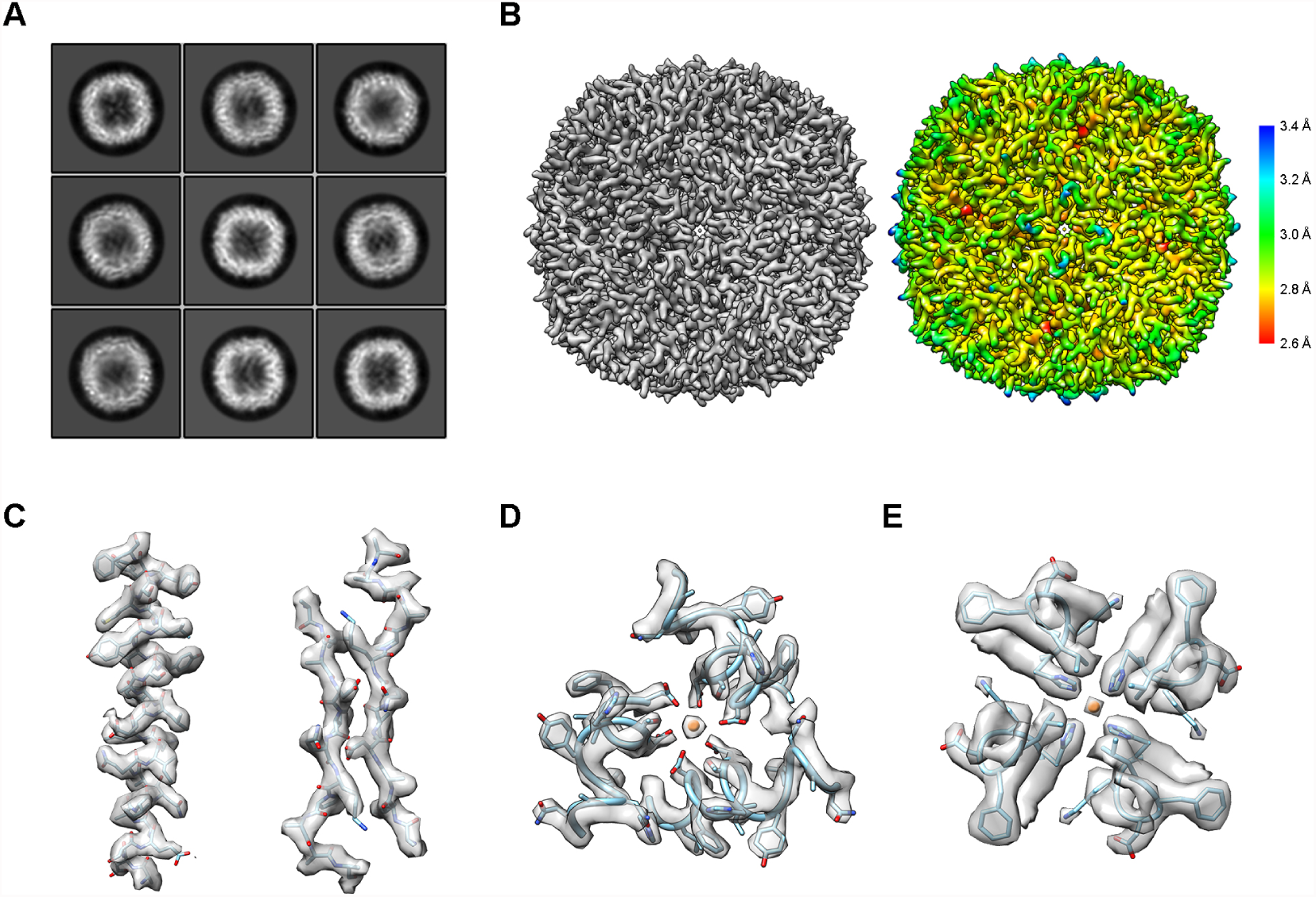
3D reconstruction of mixed under/over-focused dataset of apo-ferritin. (A) Representative 2D class averages. (B) The 3D reconstruction map at 3.0 Angstrom resolution (left) and the local resolution map (right). (C) Representative densities of secondary structures: α-helix (left) and β-sheet (right). (D and E) Ion binding sites of apo-ferritin at its 3-fold (D) and 4-fold (E) symmetric axis.

We evaluated the cryo-EM reconstructions and compared them with previously solved crystal structures of apo-ferritin. Using the reconstruction from the mixed under/over-focused dataset as an example, as shown in the local resolution map, the reconstruction as a hollow spherical assembly has the highest resolution in the middle of the shell, reaching to 2.6 Å resolution, and low resolution on the inner and outer surfaces of the assembly (Figure 2B). The quality of the map is good enough to dock a crystal structure of apo-ferrtin (PDB: 1FHA) (Lawson and Smith, 1991) into the density map with high correlation (0.9078). All the key features of the secondary structural elements and major side chains are clearly visible and match well (Figure 2C-2E and S2D). We then performed a refinement of the docked atomic model against the map, leading to a model with almost no difference from the crystal structure (RMSD 0.08 Å). We noticed that at this resolution of the reconstruction, the major side chains can be easily recognized, especially the aromatic amino acids and positively charged amino acids (Figure S2D). Furthermore, we can distinguish metal ions bound to both the 3- fold and 4-fold symmetry axis of the complex (Figure 2D and 2E). These proved the capability of VPP-Cs-corrector coupled cryo-EM in resolving high resolution structures using this novel under- and over-focus imaging method.

Additionally, to evaluate the quality of the raw VPP data that contributed to the final high resolution structure, we investigated the statistics of the particle distribution in the best reconstruction. As expected, particle images with phase shift ranging from 0.4-0.6π were the majority by both the absolute number and percentage for the final high resolution reconstruction, probably because of their high contrast for more accurate alignment (Figure 3A). The defocus value did not seem to have much effect on the percentage of particles contributing to the final high resolution reconstruction except that the images with absolute defocus values lower than 0.2 μm contributed less (Figure 3B). This may reflect the inaccuracy of CTF parameter determination of the near-focus images.

**Figure 3.**
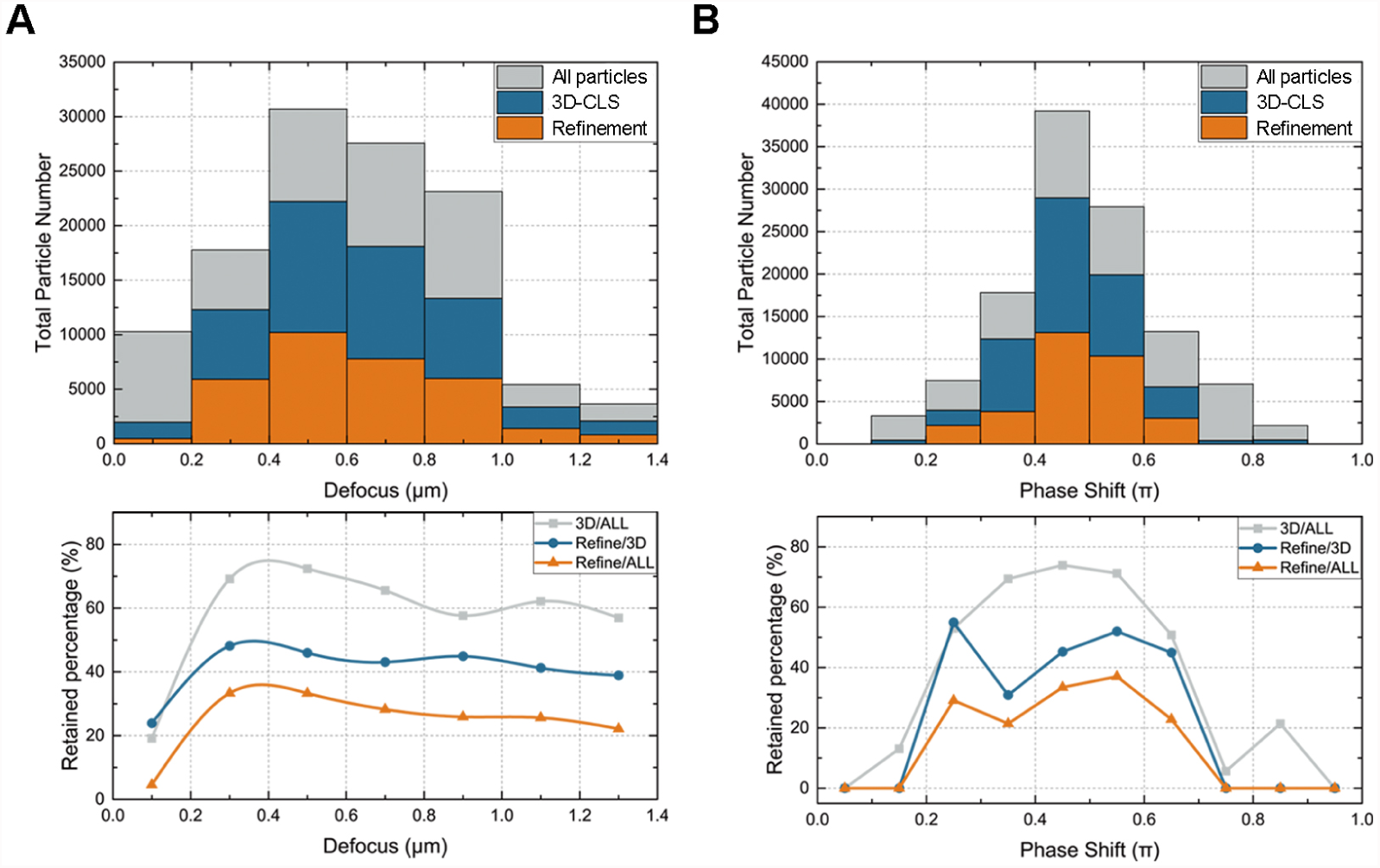
Statistics of VPP apo-ferritin reconstructions. (A) Particles distribution in different reconstruction steps according to their defocus values. In the upper panel, “All particles” (gray) contains all the particles automatically picked from the micrographs; “3D-CLS” (Blue) contains all the particles from the selected good classes after 3D classification in Relion 2.0; “Refinement” (orange) contains the particles used for the best reconstruction. The normalized ratio of retained particles in each step is in the bottom panel, where “ALL”, “3D” and “Refine” refer to “ALL particles”, “3D-CLS” and “Refinement”, respectively. (B) Particle distribution (upper) and retained percentage (underneath) according to their phase shift. The labels are the same as in (A).

### Dose-dependent reconstruction of the VPP dataset

The signal to noise ratio (SNR) of molecule images is a crucial factor for data processing. For CTEM data collection, usually high electron dose or high defocus are used to increase the SNR, especially at the low frequency. But the high electron dose would cause radiation damage of the specimen and the high defocus value would cause inaccuracy of CTF correction at high frequency, both dampening the high frequency signal. We analyzed the dose-dependence of reconstruction from the VPP dataset. When using CTEM without VPP, in order to clearly see the protein particles, we generally use a total dose 40 to 50 e^-^/Å^2^ and defocus values ranging from −1.0 to −2.0 μm to collect movie stacks. In our case, VPP movie stacks of apo-ferritin were collected with 25 e^-^/Å^2^ total dose and absolute defocus values ranging from 0.2 to 0.8 μm were enough for image processing (Figure 2 and S3). In order to evaluate the minimum dose required for high resolution reconstruction by VPP cryo-EM, we generated summed images with different frame numbers from the original 33-frame movie stacks (Figure 4). In our dataset, we found that a total dose of ~15 e^-^/Å^2^ is sufficient for successful global angular search to reach 3.1 Å resolution (Figure 4A). Following a successful global angular search, local search and refinement can be done with only the first 10 frames (a total dose of 7 e^-^/Å^2^) to reach high enough resolution to resolve the fine details (Figure 4B, 4C). As a matter of fact, the first 10 frames maintained the high frequency signal better than more frames summed (Figure 4C), demonstrating the accumulation of radiation damage during the exposure process.

**Figure 4.**
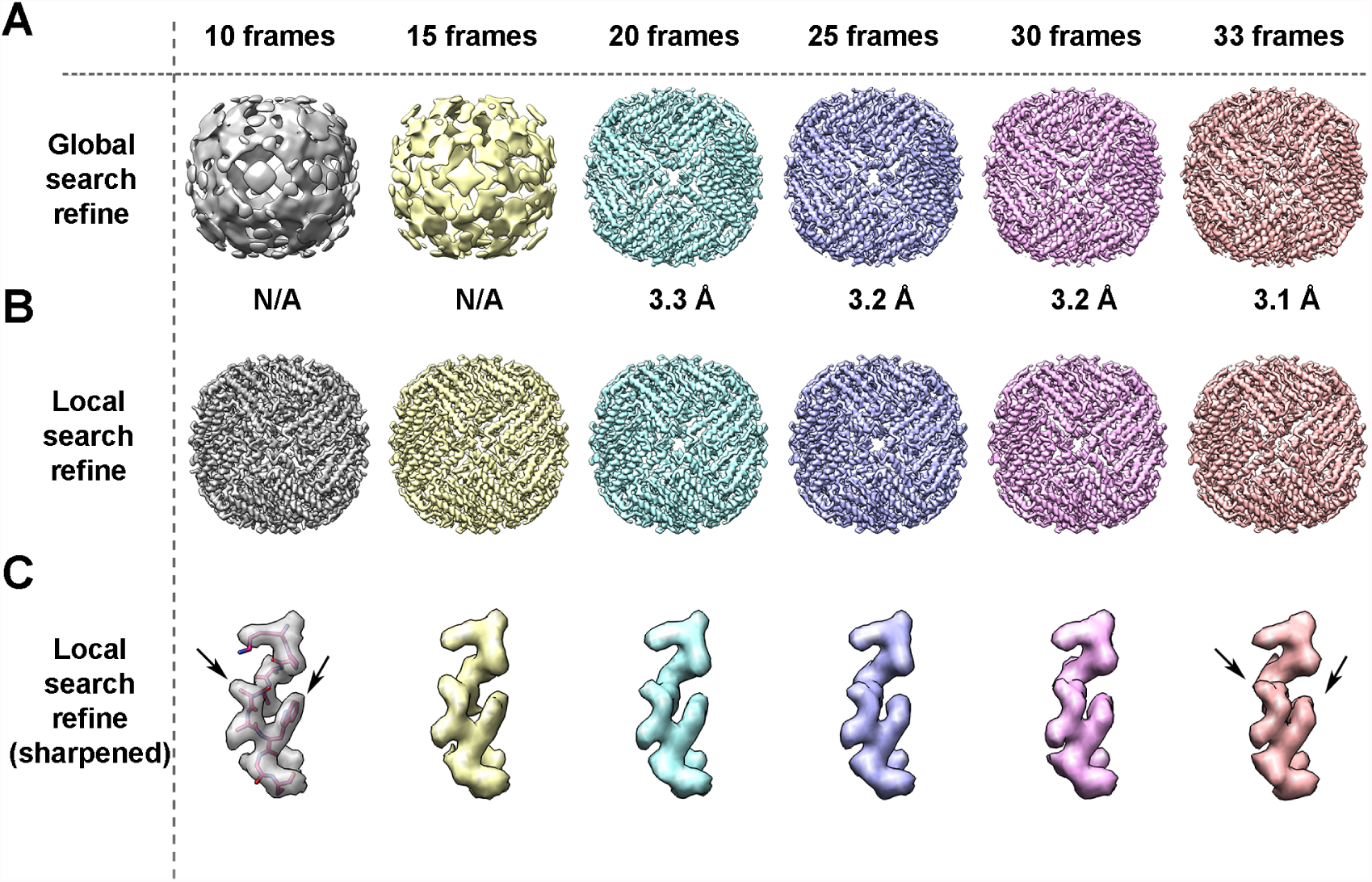
Dose-dependent reconstructions. (A) The global search reconstructions resulted from different frame combinations. The first 10, 15, 20, 25, 30 frames (7.6, 11.4, 15.2, 19, 22.4 e^-^/Å^2^ in total dose) and full 33 frames (25 e^-^/Å^2^ in total dose) are motion-corrected, summed, and refined using global angular search in Relion 2.0 individually with the same setting (7.5 degree initial angular sampling with 5 pixels initial offset range). Unsharpened maps (without post-process) are shown here with the reported resolution labeled underneath. “N/A” means that the reconstruction didn’t reach correct convergence. (B) The local-search refinement results from different frame combinations. The angular and offset information from the best reconstruction are applied for a local search from 1.8 degree with 3 pixels offset range. All the reconstructions reached to very similar resolution. (C) Representative region (loop with residue 87-95) from the sharpened maps of apoferritin (B-factor −150 Å^−2^) from each local search refinement. The corresponding atomic model was fit in the first map. Black arrows indicate the density deteriorated with more dose accumulation.

## Discussion

Cryo-EM has led structural biology into a new era, in which atomic or near-atomic resolution of macromolecule structures are much easier to be acquired (Bai et al., 2015, Nogales and Scheres, 2015). Researches have demonstrated that the application of phase plate or Cs-corrected cryo-EM respectively can solve high resolution structures (Danev and Baumeister, 2016, Fischer et al., 2015, Chua et al., 2016, Danev et al., 2017, Khoshouei et al., 2016). To our knowledge, until now there has been no high-resolution structure reported using a combination of the phase plate and Cs-corrector or determined from data collected in over-focus imaging conditions. The case study of apo-ferritin in this work proved in principle and set a novel data acquisition strategy for atomic resolution structure determination using the VPP-Cs-corrector coupled cryo-EM.

The successful reconstruction of apo-ferritin at atomic resolution in over-focus imaging condition by VPP-Cs-corrector coupled cryo-EM not only proves the feasibility of such a combined optical system, but also introduces some interesting advantages of the system for the cryo-EM community to further investigate. Since the under- and over-focused images are indiscriminate in this system, we could use a data acquisition strategy by approximately adjusting the specimen to eucentric height with the objective lens set at eucentric focus and then automatically collecting data in nearby areas by moving Compustage-XY without further adjusting the objective lens for focusing. This strategy, which keeps the stable optical system in EM during collection, may significantly speed up the data collection and reduce the lens-dependent astigmatism (Yan et al., 2017) during automatic data collection, thus improving the data quality and quantity simultaneously. This system would also change tomography data collection strategy. Currently, tomography data collection requires the whole tilting series in the under-focus imaging condition for data processing and thus causes a higher defocus values for images collected at high tilt angles. If using the VPP-Cs-corrector coupled system, one may collect tilt series at much lower defocus values, which introduces less phase inversions in high frequency thus increases the fidelity of CTF correction for tomography reconstruction. It is also intriguing to find that the minimum total dose used for high resolution refinement on the VPP-Cs-corrector coupled microscope can be as low as 7 e^-^/Å^2^. This illustrates the power of the new optical systems in cryo-EM of biological specimens, especially for cryo-electron tomography where multiple exposures at very low dose are favored.

Besides the data acquisition property, the combination of Cs-corrector and VPP also shows a better optical property than being used separately. In a simulation, we found that a relatively large astigmatism could make the two-dimensional CTF pattern distorted from oval to hyperbolic shape at high frequency area in a system with Cs (Figure S3A). This would be challenging for both CTF estimation and correction, especially for near-focus images. Such a behavior caused by astigmatism and Cs are more devastating for VPP images, because, as a matter of practical fact, VPP may introduce extra astigmatism to final images. In a system with Cs-corrector that reduces the Cs value, the hyperbolic shape disappears from the astigmatic CTF pattern therefore high frequency signals are well retained (Figure S3B).

The optical system of Cs-corrector coupled with VPP needs a stable alignment of the Cs-corrector and VPP during the image acquisition. We therefore tried to minimize the adjustment of the objective lens current (defocus value) and the retraction and insertion operation of the VPP aperture. Instead, we maintained the system at eucentric focus and adjusted the defocus by directly changing the Z-height of the specimen with the Compustage to maintain the Cs-corrector in its best performance at the preset eucentric focus. We used a modified version of the AutoEMation (Lei and Frank, 2005) software to meet our need on the VPP-Cs-corrector coupled Titan Krios instrument for a semiautomatic data collection. This software can be further improved to become fully automatic in the future, especially given that focusing step could be largely eliminated due to the system’s novel property. In the future microscope, when the next generation of Cs-corrector becomes more stable and is able to correct off-axis coma, and the phase plate becomes more robust, the combination of phase plate and Cs-corrector may become a powerful tool for high resolution cryo-EM.

In conclusion, with this work we have demonstrated the feasibility of getting high-resolution structures in over-focus imaging condition with the combination of Volta phase plate and Cs-corrector, potentiating the use of this method as a new approach for cryo-EM data acquisition. Further experiments with smaller or low-symmetry samples should be performed in the future and the advantages introduced by this novel system should be further explored for electron tomography.

## Author Contributions

X.F., L.Z., J.L., and H.W. conceived the experiments and wrote/corrected the manuscript. X.F. and L.Z. performed the cryo-EM sample preparation, data collection and processing. J.Z. and H.P. prepared graphene grids. C.L. aided in grid preparation. K.F. and X.Y. provided apo-ferritin sample. J.L. aided in hardware setup and data collection.

## Acknowledgement

Our sincere thanks to R.M. Glaser (UC-Berkeley) and R. Danev (Max-Plank Insitute of Biochemistry) for their helpful advice in setting up the VPP. We thank Y. Deng (FEI Company) and K. Zhang (MRC-LMB) for the helpful discussions of CTF fitting and correction, Xiaomin Li and Tao Yang at the Tsinghua University Branch of the National Protein Science Facility (Beijing) for their technical support on the Cryo-EM and High-Performance Computation platforms, and Fei Sun (Institute of Biophysics, CAS) for his help for the apo-ferritin sample. This work was supported by grant (2016YFA0501100 to H.W.) from the Ministry of Science and Technology of China, grant (Z161100000116034 to H.W.) from the Beijing Municipal Science & Technology Commission. K. Fan is supported by the Young Elite Scientist Sponsorship Program by CAST, Beijing Natural Science Foundation (No. 5164037), China Postdoctoral Science Foundation (No. 2015M570158) and the China Postdoctoral Science Special Foundation (No. 2016T90143).

